# Global Structural Brain-Age in Adolescence: Application of an Established Method to Short-Interval, Longitudinal Data

**DOI:** 10.64898/2026.07.31.741967

**Authors:** Amanda Boyes, Laura K M Han, Maddison Crethar, Timothy J Silk, Nandita Vijayakumar, Daniel F Hermens

## Abstract

Brain-age estimates from grey matter structure provide a promising tool to examine the neurobiology of mental health. However, whether existing models generalise to longitudinal adolescent data remains unclear. The CentileBrain Global-BrainAGE ‘Lifespan’ Model, was applied to *N*=138 participants (74 females, 64 males) from the Longitudinal Adolescent Brain Study. Participants completed 2-14 MRI scans between the ages of 12-17 years (752 datapoints). Model fit, prediction accuracy and longitudinal consistency were examined. Results showed moderate-to-good longitudinal consistency and reliability, consistent with high-performing cross-sectional age-to-brain-age correlations in youth cohorts. However, the model systematically overestimated brain-age changes relative to chronological changes, and prediction accuracy was unstable, with the mean absolute error increasing with age. Despite sex-specific brain-age calculations, on average, females showed older brain-ages compared with males. Further, younger adolescents were underpredicted, while older adolescents were overpredicted, indicating that standard model adjustments and assumptions may not be applicable.

## 1. Introduction

Adolescence is a dynamic period of neural development, coinciding with the emergence of psychiatric disorders (1). Neuroimaging research has identified distinct developmental trajectories of cortical and subcortical structure associated with the adolescent period (2, 3). While overall grey matter volume declines and white matter volume increases, different spatial, temporal and sex-specific patterns have been observed (4). Cortical grey matter volume declines non-linearly, with reductions in cortical thickness and surface area, while subcortical regions appear to peak in volume during adolescence, before declining (3, 5). In recent years, machine-learning methods have enabled the detection of biological patterns to quantify aging and interpret an individual’s ‘biological age’ (as distinct from chronological age) (6). ‘Brain-age’ models have emerged to detect deviations from normative development, by combining a range of structural features (such as thickness, surface area and volume) to determine an individual’s neurobiological age (7, 8). These models suggest that an individual’s predicted brain-age may differ to their chronological age, and recent research has examined potential influences on brain aging, across biopsychosocial, psychiatric and environmental factors (9-11).

The ‘brain-age gap’ is derived by subtracting chronological age from the estimated biological age (8). A wide gap between predicted brain-age and chronological age suggests deviation from normative developmental trajectories (5). However, brain-age models have reported an age bias in adult cohorts, where younger subjects are predicted to be older, and older subjects are predicted to be younger, indicating regression towards the mean (12).

Further, interpretation of brain-age in child-adolescent cohorts differs to adults. A higher brainage is interpreted as advanced maturation in childhood and adolescence and decline in adulthood, while a lower brain-age can suggest delayed maturation in young people and improved brain health in adulthood (13). There is also evidence of individual and context-specific variability in childhood-adolescent brain development that adds complexity to determining which brain-age deviations may be atypical (14, 15). Consequently, the degree of typical variability, compared with accelerated or decelerated brain-aging in adolescence has not yet been determined (13).

Deviations from expected trajectories of brain morphology may suggest presence or risk of disease or illness (5, 9, 16). Exploration of mental health associations with predicted brain-age and brain-age gap to date has primarily involved adult cohorts and case-control, cross-sectional studies (8, 17). Positive brain-age gap has been associated with higher mortality risk, childhood trauma, health factors (such as higher BMI, smoker status and education level), as well as depression and risk of Alzheimer’s disease in adults (17-20). Associations between brain-age and mental health may be more complex in younger cohorts. Accelerated brain aging (positive brain-age gap) has been associated with incidence of depression, externalising problems, first-episode mania, as well as psychosis, obsessive-compulsive and general psychopathology symptoms in adolescents (5, 10, 11, 21). Conversely, other research has found no association between brain-age and mental health problems in adolescents, variability in the strength of associations with different subsets of structural features, puberty-related sex differences, and associations between negative brain-age gaps and poorer mental health (13, 22, 23). There is limited evidence exploring how psychosocial and behavioural factors influence brain-age trajectories in adolescence, and a lack of longitudinal evaluation in community cohorts.

Individuals may have genetic predispositions to specific brain-age trajectories, while changes in environment may also be linked to changes in adolescent neurobiology and cognition (16, 24). Thus, while there is conceptual support for the reduction of multiple features into a single ‘score’ that captures an individual’s neurodevelopment, mental health and/or functioning, the best way to do this remains unclear.

MacSweeney et al (2025) investigated longitudinal associations between mental health and three cohort-specific measures of brain-age (25). Brain-age models were developed using white matter, grey matter and resting state functional activity in the ABCD (Adolescent Brain Cognitive Development^**sm**^) Study cohort (25, 26). White and grey matter brain-age models were more accurate (higher age correlations) than the functional activity model (25). Brain-age at baseline (mean age 9.9 years) and change in brain-age at 2-year follow-up (mean age 11.9 years) were examined with self-reported internalising problems measured at 3-year follow up (mean age 12.9 years) (25). They found that an increase in brain-age gap between timepoints (but not brain-age at baseline) was positively associated with internalising problems at 3-year follow up in females only (25). This study demonstrates sex-specific patterns and the utility of longitudinal data within a restricted age range (as data was limited to young people aged 9-11 with yearly assessment intervals). Given that adolescence represents a unique period of substantial neurobiological change and the typical age of onset for many mental health conditions (27), further research examining brain-age models in adolescent cohorts is warranted. To advance understanding of brain-age trajectories during adolescence and inform methodological considerations, including which models (algorithms, and type of imaging data) to use, how to interpret and correct bias, sample size requirements and make sense of results, short-interval longitudinal data is needed (13, 28).

Recently published research has investigated the performance of five brain-age models (trained on adult data), in the context of two cross-sectional youth cohorts (28). The study builds on previous work that has explored methodological questions such as the impact of data quality on brain-age estimates (29), and how precise or variable models should be, in order to meaningfully examine relationships with mental health (30). Three algorithms (Drobinin, Pyment, and Centile) emerged as having strong correlations with age (.51-.68) in the Michigan Twin Neurogenetics Study (MTwiNS) study, which included N=593 participants aged 9-19 years (28). However, there were only weak associations between brain-age and age in the Study of Adolescent Neural Development (SAND) cohort of N=198 Black low-income youth aged 15-17 years, with follow-up analyses indicating that the likely source of poor prediction in the SAND cohort was the restricted age (28). Thus, the authors argued that there is a need to further examine these brain-age models: (i) with longitudinal adolescent data at short intervals; (ii) in samples that overlap in age range to the training data for the model, and (iii) across wider age ranges.

The state of current research highlights that as a field, we are still understanding how to make sense of brain-age models and how to apply brain-age metrics across varied cohorts to enable meaningful conclusions. For these models to be applicable in real-world settings, they should generalise well to new datasets, produce reliable predicted ages across repeated measurements, and demonstrate longitudinal consistency whereby predicted age typically increases over time (31). Thus, the aim of the current, exploratory study was to examine the applicability of the CentileBrain Global-BrainAGE ‘Lifespan’ Model (7) to short-interval adolescent data.

## 2. Method

### 2.1 Study design and participants

The current study utilised structural neuroimaging (MRI) and demographic (age, sex) data for participants who completed at least two timepoints in the Longitudinal Adolescent Brain Study (LABS) (32) between July 2018-February 2025. Ethics approval was granted by the UniSC Human Research Ethics Committee (A181064) and written consent was obtained from all participants and parents/caregivers. LABS is ‘temporally rich’, as it recruits young people aged 12-15 years (proficient in spoken and written English) and follows them up every four months until age 17 (i.e., up to 15 timepoints (TP)). Young people are excluded from entering the study if they have suffered a major neurological disorder or major medical illness, have an intellectual disability, sustained a head injury (with loss of consciousness for more than 30 minutes) or if they are not able to complete an MRI. Participants (N=138, 74 females, 64 males; 752 datapoints) were aged 12 years (M=12.6, SD=0.3) at TP1 and 17 years (M=17.4, SD=0.25) at TP15. For sample detail by TP see *Supplementary Materials S1.1*.

### 2.2 Magnetic Resonance Imaging acquisition, processing and quality assessment

The current study utilised brain structural data from T1-weighted magnetization prepared rapid acquisition gradient echo sequence (MPRAGE; TR=2200ms, TE=1.71ms, TI=850ms, flip angle=7°, spatial resolution=0.9x0.89x0.89mm, FOV=208x256x256, TA=3:57). Scans were acquired at the Nola Thompson Centre for Advanced Imaging (Thompson Institute, UniSC) using a 3-Tesla Siemens Skyra scanner (Erlangen, Germany) with a 64-channel head/neck coil. For more detail on processing and QC, see *Supplementary Materials S1.2*.

### 2.3 Brain-age prediction model

CentileBrain Global-BrainAGE ‘Lifespan’ Model, is an open-science, web-based platform that utilises 150 FreeSurfer-processed morphometric features (68 cortical thickness, 68 cortical surface area and 14 subcortical volumes). The model has been updated in line with recent research, which found performance improved when training data was divided into age ‘bins’ 5-40 years and 40-90 years (33). The 5≤age≤40 years model (*version at September 2025*) provided sex-specific estimations of brain-age and brain-age gap. For more detail on the model and these measures, see *Supplementary Materials S1.3*.

### 2.4 Statistical analyses

The model-fit between brain-age as calculated by the established CentileBrain Global-BrainAGE ‘Lifespan’ Model and the temporally rich LABS dataset, was investigated through:(i)descriptives, (ii) visual inspection of plots, and (iii) statistical analyses. Brain-age and brain-age gap data was examined to determine: (i) model bias (for example, regression to the mean),(i)prediction accuracy, and (iii) longitudinal consistency. Data was examined using SPSS Statistics Version 29.0.0.0 (241) and RStudio (version 2024.04.2+764), “Chocolate Cosmos” Release for Windows.

### 2.4.1 Descriptives and plots

Plots were generated to visually examine: (i) the relationship between chronological age and both brain-age and brain-age gap, (ii) the residuals and mean absolute error of brain-age across polynomial age models, and (iii) change in brain-age and change in chronological over ages 12-17.

### 2.4.2 Model fit: Model residuals and ANOVA analyses

To examine model fit/bias, the model residuals of predicted brain-age were compared across four increasingly flexible polynomial mixed-effects models: (i) linear age, (ii) linear + quadratic age, (iii) linear + quadratic + cubic age, and (iv) linear + quadratic + cubic + quartic age. An ANOVA was performed to compare the results of each model and determine whether model errors were randomly distributed across age or whether systematic biases emerged at specific developmental periods.

### 2.4.3 Prediction accuracy: Mean absolute error

The mean absolute error (MAE) was computed for each age year (or ‘age bin’) across ages 12-17, along with standard errors, to determine which polynomial model showed the best prediction accuracy.

### 2.4.4 Longitudinal consistency and reliability

The longitudinal consistency and reliability of the model was assessed by: (i) calculating correlations between changes in brain-age and chronological age, between first and last timepoints for all N=138 participants. Outliers (by timepoint) were identified using the interquartile range (IQR) method and removed for both the brain-age and chronological age data, to test if the model fit improved; and (ii) Intraclass Correlation Coefficient (ICC) estimates between all pairs of consecutive timepoints (only participants who contributed to both timepoints were included in each ICC).

### 2.4.5 Sensitivity analyses: Evaluation of age range of best fit

Given the increased variability in MAE observed in the older age bins (see *Results 3.2*), model performance across the whole LABS dataset (12-17.9 years) was compared with two restricted age ranges: (i) 12-16.9 years, and (ii) 12-15.9 years. Comparisons were made on: (i) correlations between change in brain-age and change in chronological age (see 2.4.4 (i)); and (ii) linear mixed-effects models examining the association between chronological age and predicted brain-age, with subject ID included as a random intercept to account for repeated measurements.

## 3. Results

### 3.1 Descriptives

Visual inspection of the relationship between chronological age and brain-age revealed that the latter is underpredicted in younger adolescents (12-14 years) and over-predicted in older adolescents (16-17 years), in both males and females (see Figures S2 and S3). Further, the slope increases for older adolescents. Females had older brain-ages, on average, over time, when compared with males (see Figure S4).

### 3.2 Model fit and MAE

As the expected regression-to-mean bias was not observed, to determine whether the observed age bias could be reduced and model fit improved, model residuals and ANOVA analyses compared four progressively complex polynomial models, including linear, quadratic, cubic, and quartic age terms (see Figure S5 and Table S3). Results indicated that adding age^2^ significantly improved the linear age only model, and adding age^3^ further improved the model, but adding age^4^ did not. This suggests the best fit was model (iii) including linear + quadratic + cubic age. The MAE across the models was similar, increasing with older age. Figure S6 shows that higher-order polynomial terms provided minimal improvements in reducing prediction error. MAE values increased by ∼50% and became more variable over time, across all models.

### 3.3 Longitudinal consistency and reliability

Brain-age and brain-age gap, plotted by individuals over time are included in *Supplementary Materials S2.3* (Figures S7 and S8). Brain-age predictions demonstrated moderate-to-good longitudinal consistency, with an individual’s brain-age change (from their first to their last scan) significantly correlated with their chronological age change (*r*=.63, 95% CI .52-.72, *p* <.001) (Table 1). Significant departures from normality were observed for both chronological age change (Shapiro-Wilk *W* = 0.89, *p* < .001) and brain-age change (*W* = 0.97, *p* = .011). Five participants had data identified as outliers (by timepoint) for brain-age change using the IQR method. Sensitivity analyses showed that removing outliers didn’t change the results (*r* = .63, 95% CI .52-.73), thus the Pearson correlations were considered robust. Sex-stratified analyses showed comparable longitudinal consistency for males (*n* = 64, r = .60, 95% CI .41-.74, *p* < .001) and females (*n* = 74, *r* = .66, 95% CI .51-.78, *p* < .001), with no significant difference between correlations (Fisher’s *z* = −0.62, *p* = .537). However, the model systematically overestimated brain-age changes (*M* = 3.80 years, *SD* = 3.98) relative to chronological changes (*M* = 2.23 years, *SD* = 1.57), resulting in significant positive bias (mean residual = 1.57 years, 95% CI 1.02-2.11, *p* <.001) (Figure 1; Table 1). Both sexes exhibited similar systematic overestimation bias (males: *M =* 1.62 years, *SD =* 3.58; females: *M* = 1.52 years, *SD* = 2.94). Bland-Altman analysis revealed wide limits of agreement, indicating substantial individual variability in prediction accuracy (Figure S9). ICCs were ∼.55-.85 across all timepoints demonstrating moderate-to-good reliability (Table S4 and Figure S10).

**Table 1:** Longitudinal consistency across whole sample (12-17 years) and two restricted age ranges (12-15 and 12-16 years)

| Age range<br>(mean) | 12.1-15.9<br>(13.9) | 12.1-16.9<br>(14.3) | 12.1-17.9<br>(14.5) |
| --- | --- | --- | --- |
| *Descriptives* |  |  |  |
| N participants | 126 | 136 | 138 |
| Datapoints | 577 | 690 | 752 |
| Mean age at first scan | 13.04 | 13.23 | 13.27 |
| Mean age at last scan | 14.69 | 15.19 | 15.50 |
| *Chronological age* |  |  |  |
| Mean change (SD) | 1.65 (1.01) | 1.96 (1.30) | 2.23 (1.57) |
| *Brain-age* |  |  |  |
| Mean change (SD) | 2.60 (3.38) | 3.36 (3.78) | 3.80 (3.98) |
| *Inferential statistics* |  |  |  |
| Pearson $r^a$ | 0.51<sup>b</sup> | 0.62<sup>b</sup> | 0.63<sup>b</sup> |
| 95% CI lower - upper | 0.37 - 0.63 | 0.51 - 0.72 | 0.52 - 0.72 |
| R-squared | 0.26 | 0.39 | 0.40 |
| Mean bias (SD) | 0.95 (3.0) | 1.39 (3.14) | 1.57 (3.23) |

All age, change and bias scores are reported in years. <sup>a</sup>Correlation between change in chronological age and change in brain-age, between an individual's first and last scans. <sup>b</sup> $p < .001$ .

**Figure 1.**
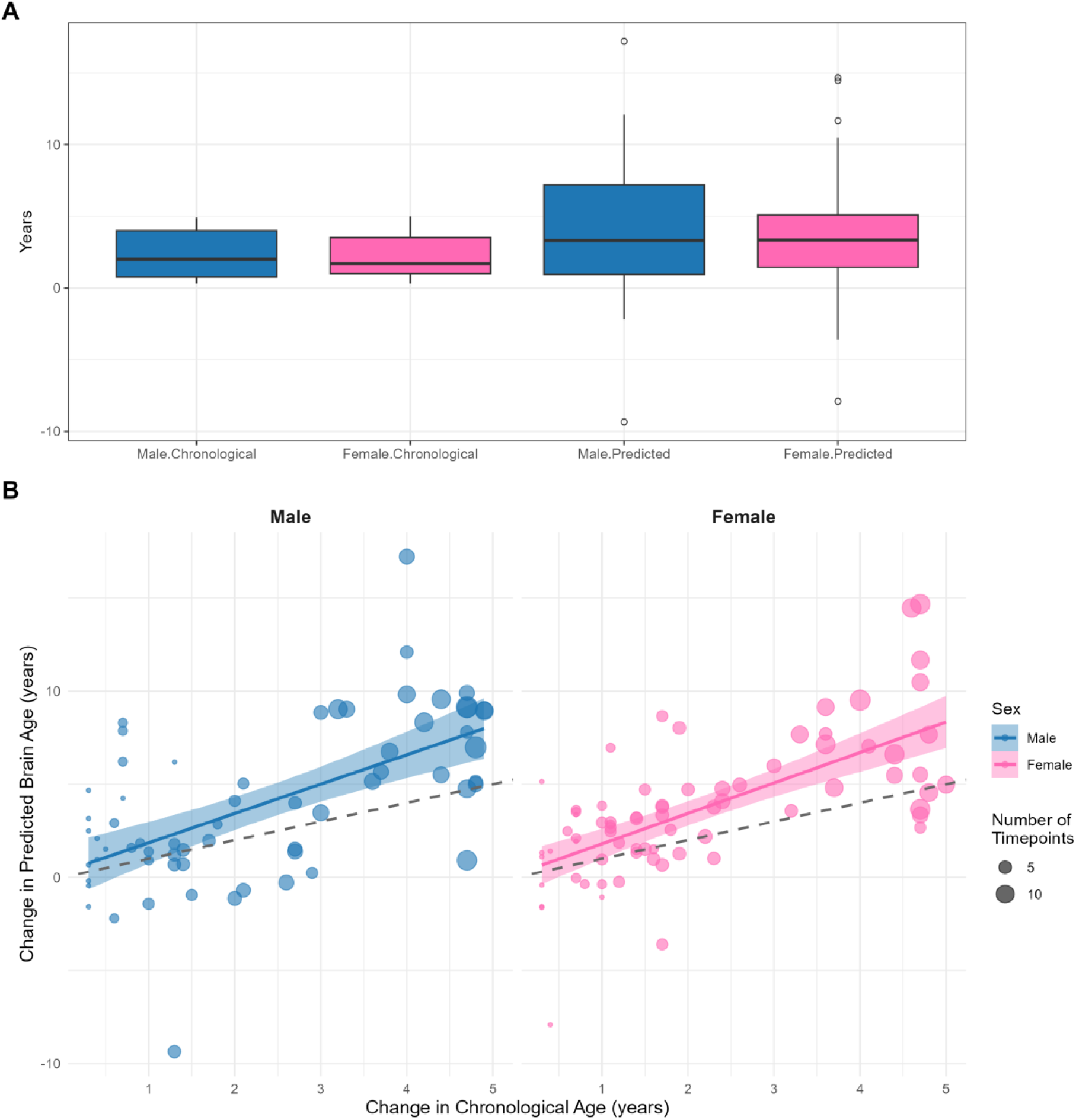
(A) Box plots for change in chronological age and change in brain-age, split by sex, across ages 12-17. (B) Change in predicted brain-age x change in chronological age, split by sex; size of dots represent number of timepoints contributed by individuals; coloured lines indicate brain-age overestimation bias with 95% confidence intervals shaded; dashed reference line indicates perfect fit.

### 3.4 Sensitivity analyses: Age range of best fit

Across the whole LABS dataset (12-17.9 years) and the two restricted age ranges (12-16.9 years, and 12-15.9 years), chronological age was positively associated with brain-age (Figures S4 and S11; Tables 1 and S5). Brain-age change (from first to last scan) in the 12-16.9 years data was similar to the whole dataset (*n =* 136, *r =* .62, 95% CI .51-.72), showing stronger correlations than the 12-15.9 years data (*n* = 126, *r* = .51, 95% CI .37-.63). All three models overestimated brain-age relative to chronological age, with bias increasing across wider age ranges (see Table 1). Variance explained increased from the 12-15.9 years model (*R*^2^ = .26) to the 12-16 model (*R*^2^ = .39), with negligible additional variance explained by the 12-17.9 years model (*R*^2^ = .40). Linear mixed-effects models examining the association between chronological age and predicted brain-age (Table S5) show that model fit (as evaluated by the REML criterion) decreases across the three models, while the age slope (fixed effect) and participant variability (random effect) increases.

## 4. Discussion

This study contributes to our understanding of the strengths and limitations of the CentileBrain BrainAGE model when applied to a temporally rich longitudinal adolescent dataset, with implications for utility in examining associations with other variables (e.g. predicting mental health outcomes), as well as further model testing and development. Firstly, change in brain-age was positively correlated with change in chronological age, and ICCs were ∼.55-.85 across all timepoints, suggesting a moderate level of prediction accuracy and reliability. However, there was an observed age bias whereby younger adolescents (12-14 years) were found to have younger brain-ages and older adolescents (16-17 years) demonstrated older brain-ages. This is opposite to what is typically observed (12, 31). Further the model systematically overestimated brain-age changes, and MAE values increased by ∼50% becoming more variable over time. ANOVAs comparing increasingly non-linear models of the residuals suggested that the best fit was achieved by combining linear + quadratic age + cubic age. Finally, despite using sex-specific templates, on average, females demonstrated older brain-ages compared with males. Taken together, this suggests that while the model performance is comparable to results observed in cross-sectional adolescent cohorts, different model adjustments may be needed when focussing on longitudinal data and/or a short developmental window within the adolescent period, compared to cross-sectional analyses, trajectories over longer timeframes and/or older cohorts. This may reflect fundamental differences between brain development and brain aging trajectories, or may be due to systematic differences between the training dataset and the LABS cohort (34). The CentileBrain BrainAGE model may not adequately capture the non-linear trajectory of grey matter across adolescent neurodevelopment (28), and increasing individual variability in adolescent brain maturation (35) may lead to greater prediction error within this developmental window.

Sensitivity analyses comparing model fit of the whole LABS dataset (12-17.9 years) with the two restricted age ranges (12-16.9 years, and 12-15.9 years) found that the 12-15.9 years data had the highest accuracy in age prediction (lowest REML, Table S5), while the 12-16.9 and 12-17.9 datasets explained the most variance (highest R^2^, Table 1). Further, the ICCs (Table 4) show that while age is tightly constrained across the timepoints and there is moderate-to-good longitudinal reliability, brain-age gap varies over time, and Table 1 shows increased heterogeneity and bias in the 12-17.9 years data. This may be the result of a small sample size across a wider age range, as there was only an additional 60 datapoints in the 17-17.9 age range and the average age and sample *n* was similar to the 12-16.9 years model. By comparison, there were an extra 106 datapoints (and an increase of *n*=10 participants) in the 12-16.9 years group, compared with the 12-15.9 restricted age dataset. Thus, the extra heterogeneity in brain-age prediction may indicate (i) developmental differences, (ii) limitations of this sample in the context of this model, and/or (iii) the optimal age range to sample size ratio in temporally rich longitudinal cohorts.

Recent research suggests that models with lower age-prediction accuracy may be more desirable when the goal is individual disease detection (36). ‘Looser-fitting’ models would also allow for determination of an individual’s unique trajectory, which is a distinguishing feature of adolescent brain development, as well as cognitive and emotional development (35). Standard adjustments to brain-age estimates that assume regression-to-the-mean may not apply. Thus, balancing model-fit considerations (such as assessing and appropriately adjusting for model bias) while also allowing for meaningful individual variability may improve replicability in adolescent brain-age associations with other factors, such as mental-health or lifestyle. Building on the results presented in the current study, CentileBrain brain-age measures were further examined in association with mental health clusters (*see Crethar et al, submitted*). By continuing to evaluate the model fit of existing brain-age models to a range of developmental cohorts, the utility and interpretation of brain-age in the adolescent period can be more fully understood.

## Supporting information

BOYES-brainage-supplemental-materials

## Acknowledgements

Thank you to the young people who gave their time to participate in this research. Thank you also to the Thompson Institute radiographers, LABS research assistants Marcella Parker, Arline Rathjen-Duffton and Shae Rendall, the students and other staff who assisted with data collection. Pete Embleton was instrumental in successfully processing the MRI data on the UniSC HPC. Monica D Rosenberg reviewed the manuscript and provided valuable feedback. The authors acknowledge the use of Claude Sonnet 5 and Microsoft 365 Copilot (bizchat.20260701.51.4) to assist with code development, syntax refinement, and troubleshooting during the data analysis process. All analytical decisions, interpretation of results, and manuscript content were reviewed and verified by the authors, who take full responsibility for the work.

## Data availability

The datasets generated and analysed during the current study are not publicly available due the fact that they constitute an excerpt of research in progress but are available from the corresponding author on reasonable request.

## Author contribution statements

Amanda Boyes: conceptualisation, methodology, formal analysis, investigation, data curation, writing – original draft, visualization, supervision, project administration, funding acquisition; Laura K M Han: conceptualisation, methodology, writing – review and editing, visualization, supervision; Maddison Crethar: methodology, formal analysis, investigation, data curation, writing – review and editing, visualization; Timothy J Silk: writing – review and editing; Nandita Vijayakumar: writing – review and editing; Daniel Hermens: conceptualisation, methodology, writing – review and editing, project administration, resources, funding acquisition.

