## Supplementary material for "Global Structural Brain-Age in Adolescence: Application of an Established Method to Short-Interval, Longitudinal Data": BOYES-brainage-supplemental-materials

**Supplemental Material**

**S1 Methods**

**S1.1 Participants**

The final dataset for this study included *N*=138 adolescents (74 females, 64 males) with a total of 752 datapoints, across up to 14 timepoints (TP), within a 5-year period (maximum of 15 TPs) at 4-monthly intervals.

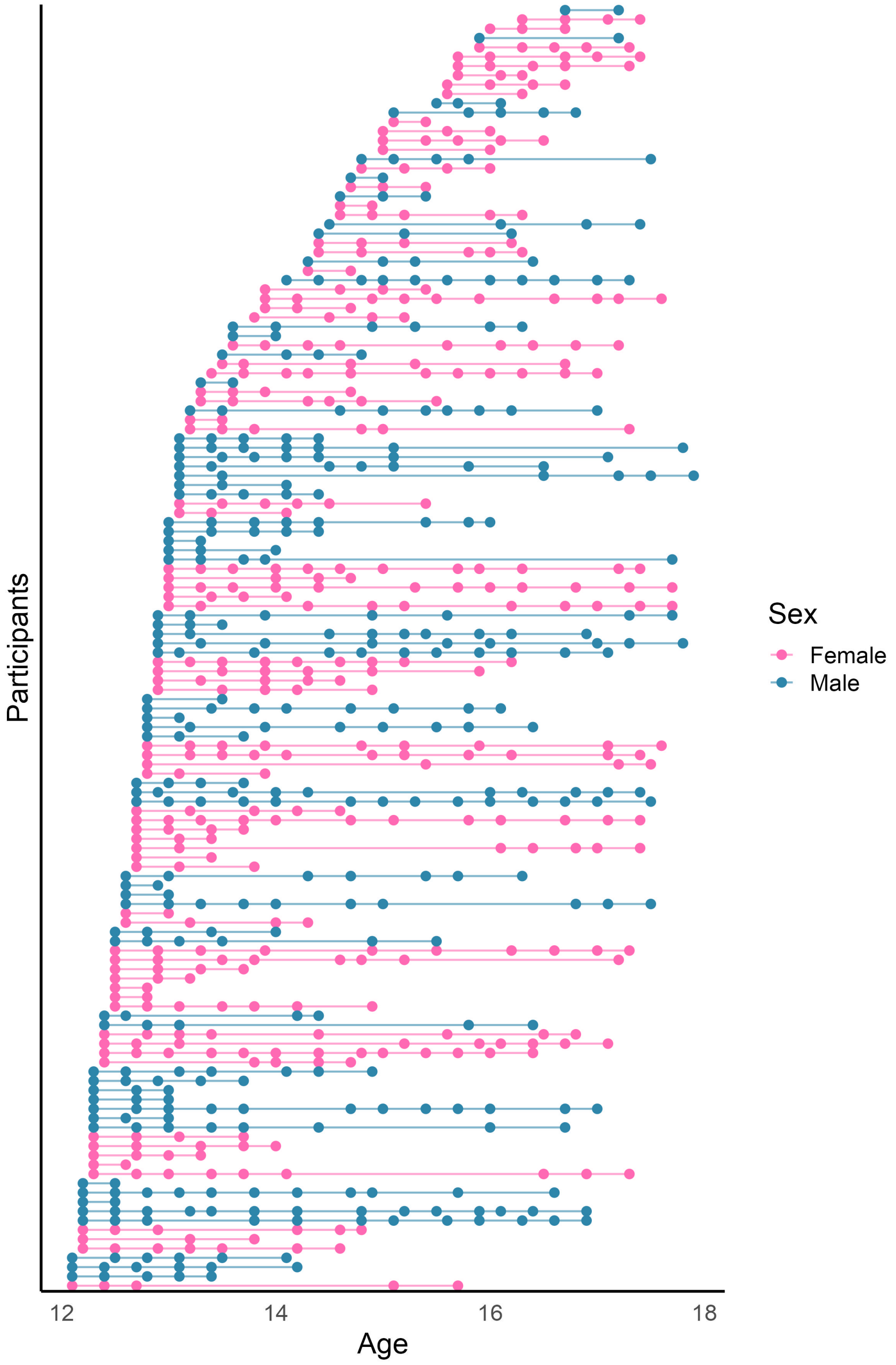


**Figure S1:** Visual representation of all data contributed to the final dataset, by individual, coded by sex.

**Table S1: Summary by timepoint: Datapoints (by sex) and age (total *N*=138; 752 datapoints)**

| Timepoint | Female datapoints (n = 74) | Male datapoints (n = 64) | Age Mean (SD) | Age Range |
|---|---|---|---|---|
| **TP1** | 39 | 42 | 12.59 (.30) | 12.1-13.0 |
| **TP2** | 36 | 45 | 12.93 (.31) | 14.4-13.5 |
| **TP3** | 34 | 33 | 13.23 (.30) | 12.7-13.9 |
| **TP4** | 32 | 32 | 13.61 (.30) | 13.1-14.2 |
| **TP5** | 34 | 26 | 13.95 (.29) | 13.4-14.5 |
| **TP6** | 20 | 16 | 14.23 (.27) | 13.8-14.8 |
| **TP7** | 29 | 28 | 14.63 (.26) | 14.1-15.1 |
| **TP8** | 29 | 21 | 14.99 (.30) | 14.5-16.5 |
| **TP9** | 20 | 20 | 15.32 (.27) | 14.8-15.8 |
| **TP10** | 21 | 17 | 15.66 (.24) | 15.1-16.0 |
| **TP11** | 24 | 18 | 16.00 (.22) | 15.5-16.5 |
| **TP12** | 19 | 15 | 16.26 (.24) | 15.7-16.8 |
| **TP13** | 19 | 14 | 16.68 (.26) | 16.1-17.2 |
| **TP14** | 19 | 16 | 16.98 (.28) | 16.3-17.5 |
| **TP15** | 18 | 16 | 17.40 (.25) | 16.9-17.9 |
| **Total** | **393** | **359** | **-** | |

**Table S2: Total number of datapoints contributed by individuals, by sex (total *N*=138; 752 datapoints)**

| Number of timepoints | Female | Male |
|---|---|---|
| 2 TPs | 11 | 12 |
| 3 TPs | 11 | 10 |
| 4 TPs | 15 | 6 |
| 5 TPs | 13 | 9 |
| 6 TPs | 5 | 5 |
| 7 TPs | 3 | 6 |
| 8 TPs | 2 | 4 |
| 9 TPs | 5 | 3 |
| 10 TPs | 3 | 2 |
| 11 TPs | 2 | 3 |
| 12 TPs | 3 | 2 |
| 13 TPs | 1 | - |
| 14 TPs | - | 2 |

As a participant may attend up to four timepoints in a 12-month period, the maximum number of timepoints (TPs) contributed by any individual to each 'age bin' was also checked and determined to be three for females and four for males (i.e., across each year of pooled data for 12, 13, 14, 15, 16, and 17 year-olds) – see Figure S3.

### S1.2 Image processing software and quality assessment

T1-weighted images were processed using the FreeSurfer (7.4.0) longitudinal recon-all pipeline (1). Software was packaged in singularityCE (version 3.8.0), on the UniSC HPC system, using PBS (version 20.0.0), Ubuntu 20.04, and connected to a CEPH cluster (version 15.2.13).

**Raw data QC**

Prior to conducting FreeSurfer analyses, scans were visually inspected and scans were removed due to either having only one scan or a poor-quality scan (i.e., movement or artefact) (n=14).

**Euler numbers**

Following FreeSurfer analyses, Euler numbers were inspected using SurfaceHoles and EstimatedTotalIntraCranialVol from the cross-sectionally-processed (CROSS) aseg FreeSurfer files. Euler numbers were calculated for all MRI scans using the below equation:

$$Euler\ number = 2 - 2\ x\ total\ FreeSurfer\ CROSS\ brain\ surface\ hole\ number$$

Euler numbers provide an indication of cortical surface topology and have been used in neurodevelopment studies as a measure of image quality (2, 3). A perfect surface with no holes would have an Euler number of two, however, in adolescent imaging studies, it has been previously reported that values can be as low as -380 and may mediate apparent relationships between grey matter and age (2).

Zscores were generated and any <-3.0 were visually inspected in FreeView and ENIGMA-generated html files.

**ENIGMA-QC**

The ENIGMA FreeSurfer Cortical and Subcortical Extraction and QC Protocol (version 2; https://enigma.ini.usc.edu/protocols/imaging-protocols/) is openly available online (https://github.com/ENIGMA-git/ENIGMA-FreeSurfer-protocol) (4). The output includes: (1) extraction of 150 cortical and subcortical measures from the FreeSurfer-processed data; (2) creation of summary statistics, flagging outliers on subcortical and cortical measures; and (3) generation of html pages with FreeSurfer images for each scan, to enable efficient QC review, including: subcortical (by region of interest), cortical (internal, all regions) and cortical (external, all regions). The ENIGMA resources include advice on how to rate each region, including reference images. Outliers on any structural metrics (i.e., thickness, surface area or subcortical volume) were reviewed.

**Final exclusions and FreeSurfer longitudinal recon-all re-processing**

Following examination of outliers identified by Euler numbers and/or the ENIGMA-QC pipeline, an additional 45 datasets (from n=7 participants) were excluded. FreeSurfer BASE and LONG brains were re-run for any participants with scans removed, prior to statistical analyses.

#### S1.3 CentileBrain Global-BrainAGE 'Lifespan' Model

The CentileBrain Global-BrainAGE 'Lifespan' 5≤age≤40 years Model (version at September 2025, *available here:* https://centilebrain.org/#/brainAge_global) included (i) *predicted age* ('brain-age' prediction, unadjusted by chronological age); (ii) *BrainAGE* ('brain-age' gap, which is the difference between chronological age and unadjusted 'brain-age' prediction); (iii) *adjusted predicted age* (the 'brain-age' prediction, adjusted for chronological age); and (iv) *adjusted BrainAGE* (adjusted brain-age' gap, which is the difference between chronological age and adjusted 'brain-age' prediction). For additional description of the model algorithm see (5). The current study examined (i) referred to as 'brain-age' and (iv) referred to as 'brain-age gap'.

The CentileBrain BrainAGE model assumes the usual over/under estimation bias across age (6). As noted on the website FAQs (https://centilebrain.org/#/faq) "**How is age adjustment implemented in the brainAGE models?**" - "BrainAGE models typically show age-dependent bias (younger individuals are overpredicted and older individuals are underpredicted). Age adjustment corrects this by calibrating predicted age to chronological age using regression functions. The CentileBrain BrainAGE models optimize accuracy by using a combination of linear and non-linear functions that are customised to each uploaded dataset by estimating the adjustment from the observed age-prediction relationship in that dataset, then applying it to generate an adjusted predicted age."

### S2 Results

#### S2.1 Descriptives and visual inspection of plots

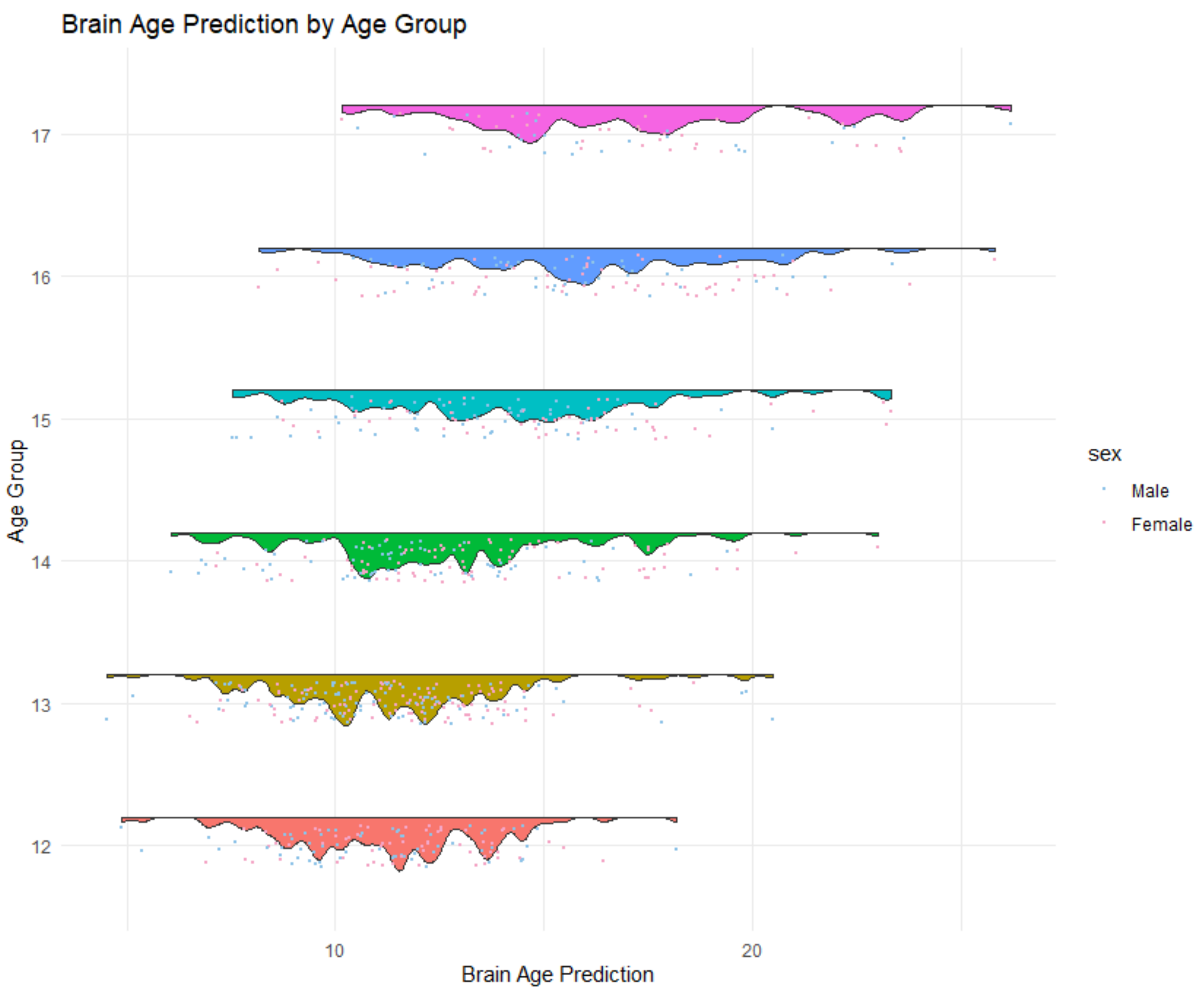


**Figure S2:** Brain-age by Chronological Age Group. Each coloured band shows the distribution of brain-age predictions for individuals in each chronological age group (e.g. 12-12.9, 13-13.9, etc). The dots represent individual data points, while the coloured regions show the density or concentration of predictions.

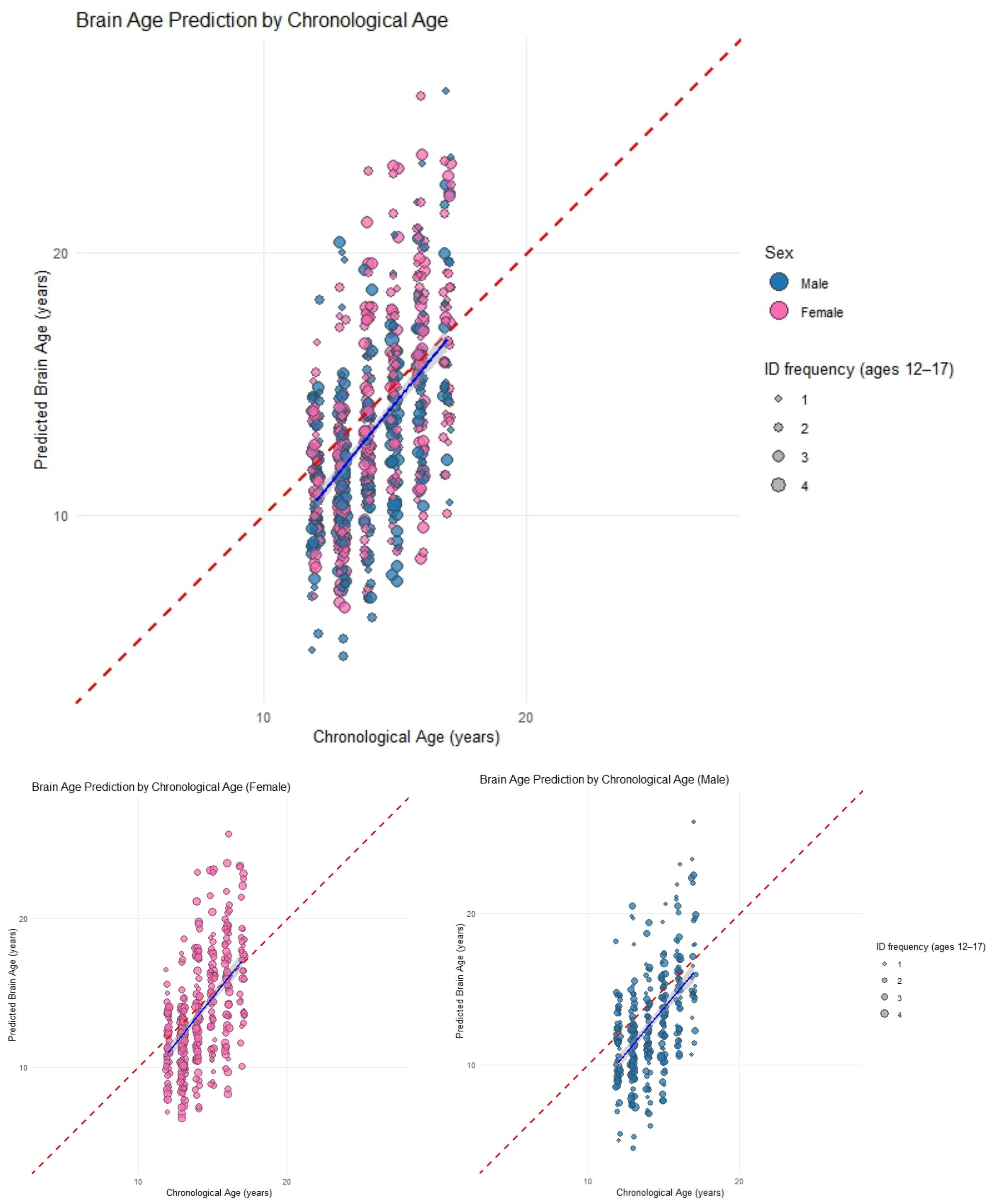


**Figure S3: Brain-age by Chronological Age x Sex.** Red dashed line indicates the perfect prediction (where brain-age = chronological age); blue lines indicate the actual relationships.

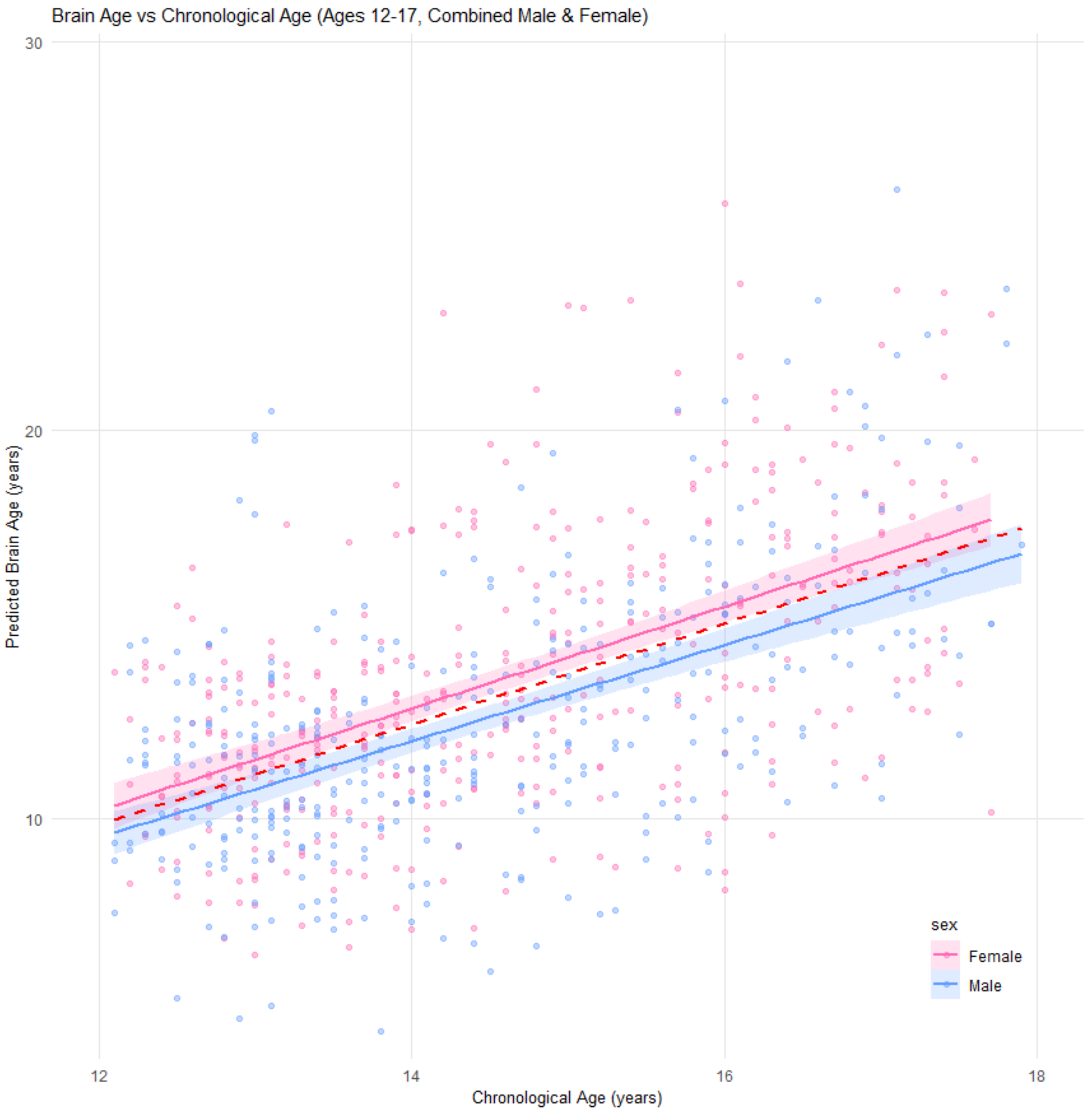


**Figure S4:** Brain-age x chronological age (12-17 years) x sex. Regression lines of best fit indicated at the group level (red dashed line), and by sex with 95% confidence intervals (shaded).

#### S2.2 Model fit and predication accuracy

#### S2.2.1 Residuals and ANOVAs

To address the observed age-bias and improve model fit, residuals (i.e., difference between brain-age and chronological age; unadjusted brain-age gap) were calculated and four models of these residuals were compared.

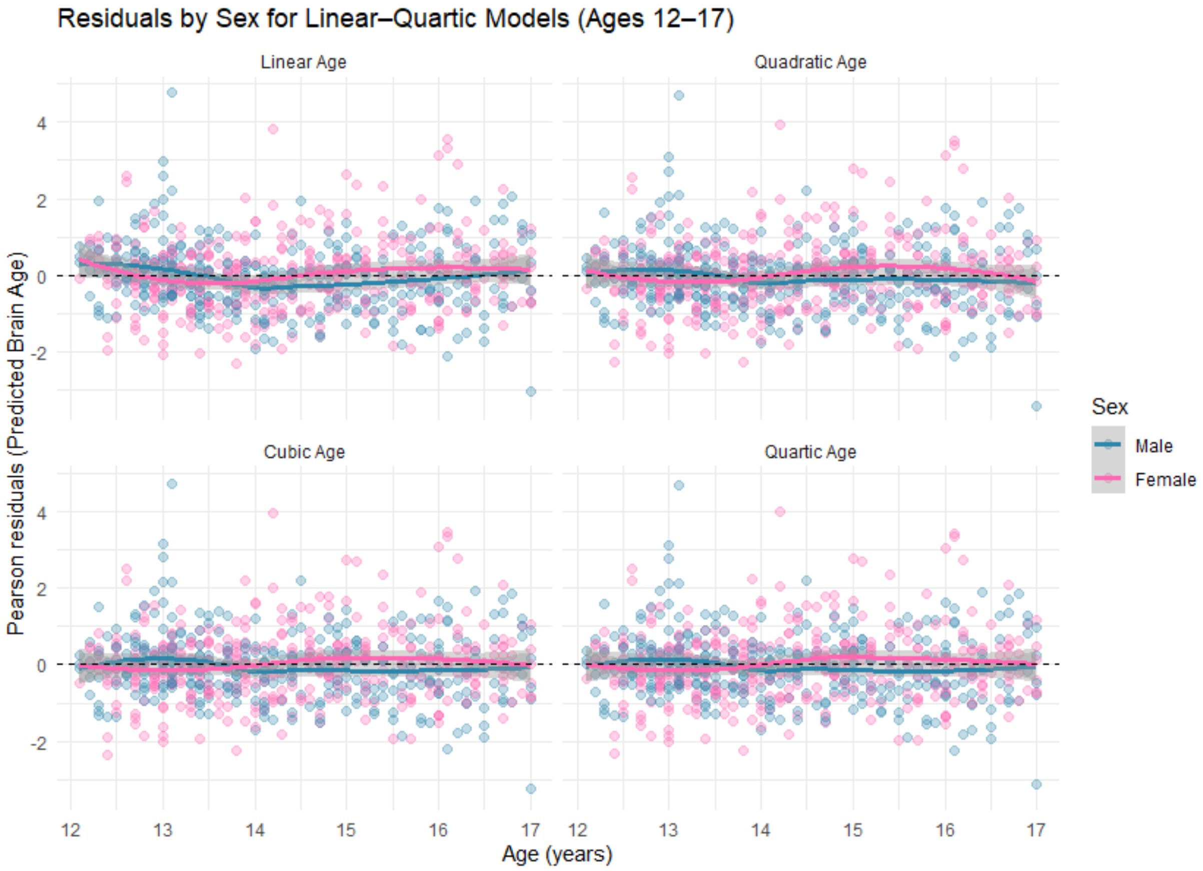


**Figure S5:** Brain-age Residuals by Sex for Linear, Quadratic, Cubic and Quartic Age Models

#### S2.2.2 ANOVA analyses

ANOVA analyses were conducted to see which model best explained the brain-age residuals (i.e., unadjusted brain-age gap).

**Table S3: ANOVA Comparison of Four Models of Brain-age Residuals**

| Model | df | AIC | BIC | logLik | Test | L.Ratio | p-value |
|---|---|---|---|---|---|---|---|
| **Model 1** (linear age) | 4 | 3400.64 | 3419.13 | -1696.32 | | | |
| **Model 2** (linear + quadratic age) | 5 | 3389.29 | 3412.40 | -1689.64 | 1 vs 2 | 13.35 | $p$ <.001 |
| **Model 3** (linear + quadratic age + cubic age) | 6 | 3386.07 | 3413.81 | -1687.03 | 2 vs 3 | 5.22 | 0.022 |
| **Model 4** (linear + quadratic age + cubic age + quartic age) | 7 | 3387.86 | 3420.22 | -1686.93 | 3 vs 4 | 0.21 | 0.645 |

#### S2.2.3 Prediction accuracy: Mean absolute error (MAE)

The MAE is the average of the absolute differences between the brain-age and observed age data (i.e., unadjusted brain-age gap). It can be used as a measure of model performance (7).

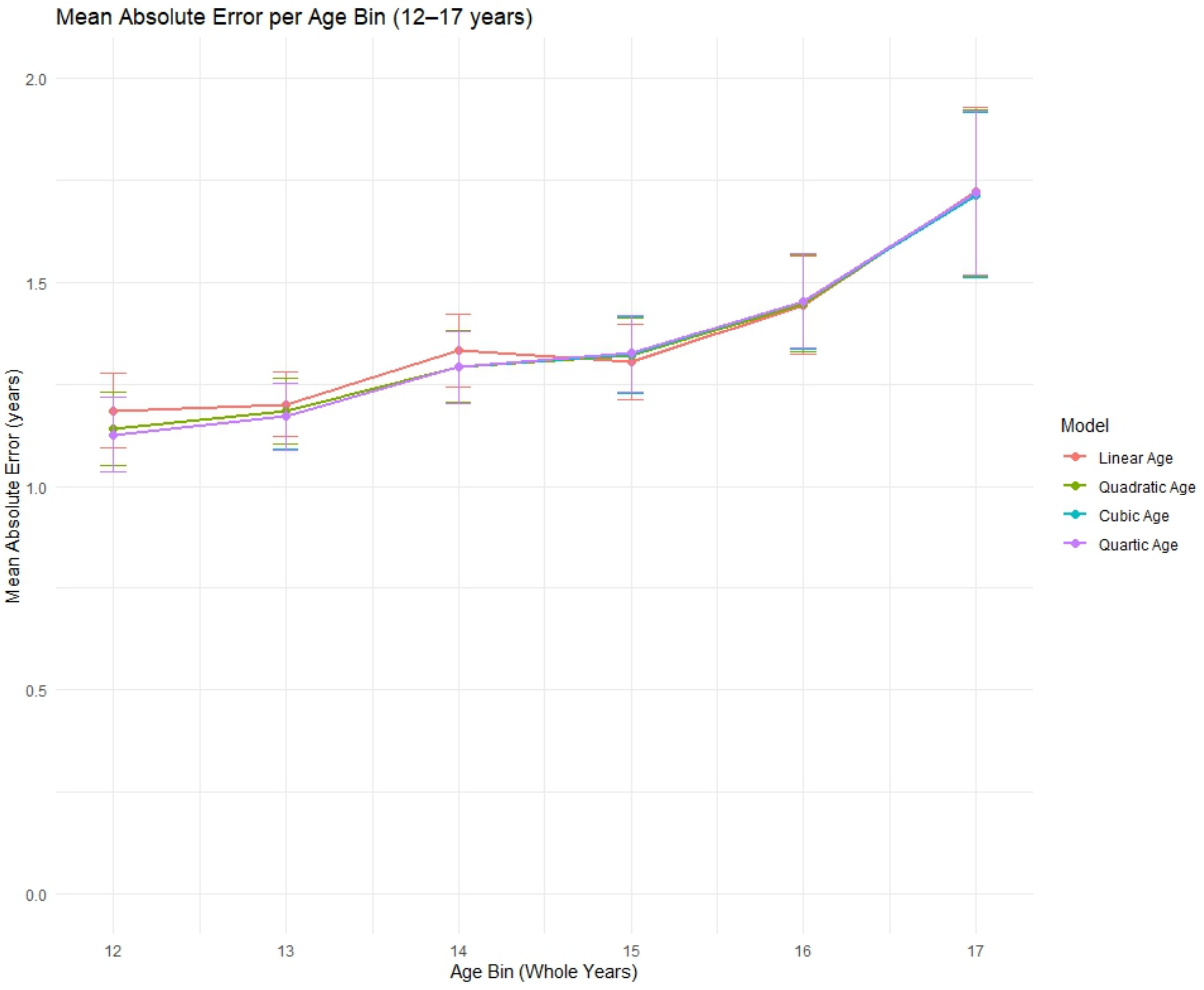


**Figure S6: Mean absolute model error per age bin**

### S2.3 Longitudinal consistency

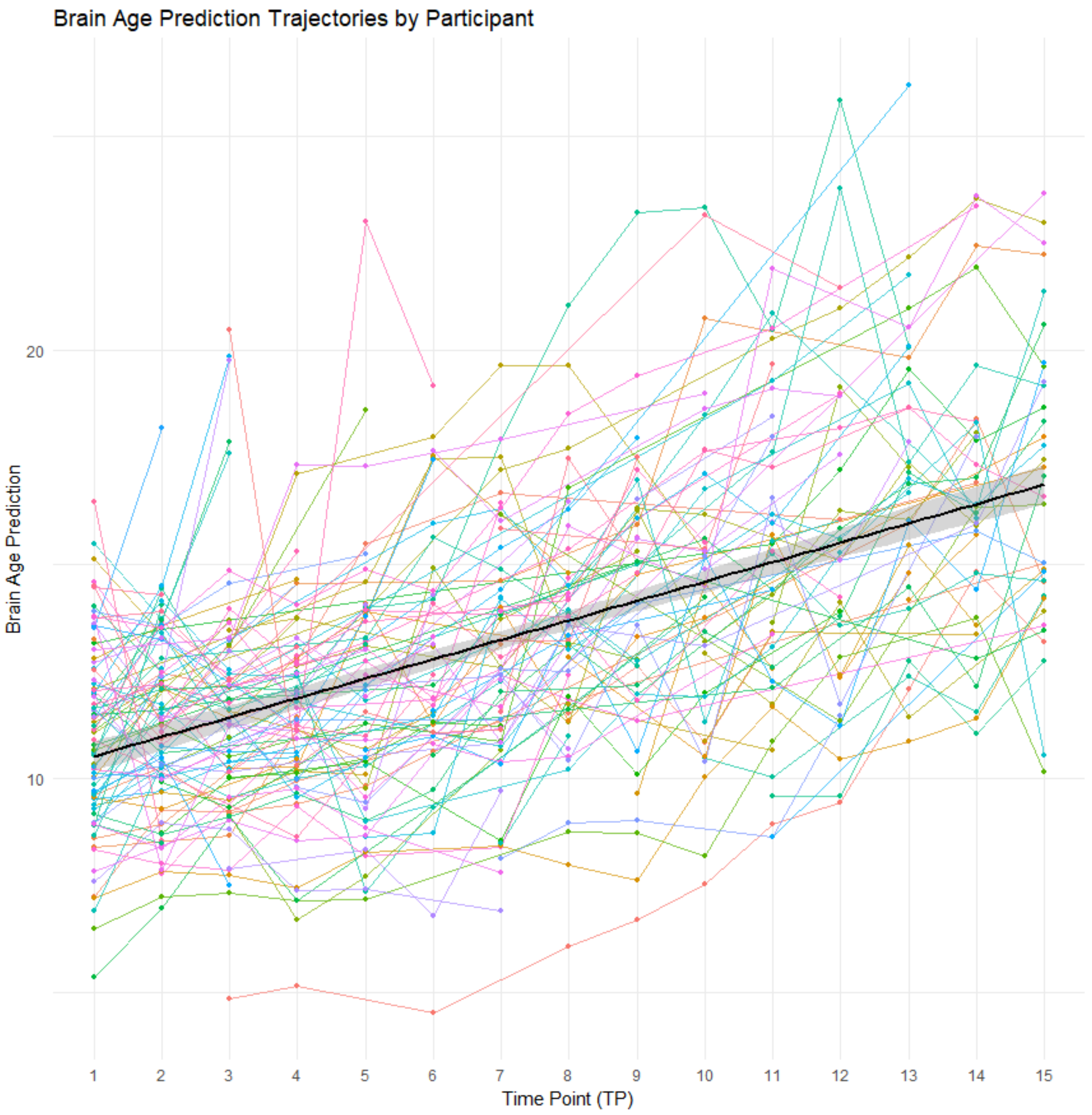


**Figure S7:** Participant Brain-age Trajectories over 12-17 years. Regression line indicates line of best fit with 95% confidence intervals (grey shading).

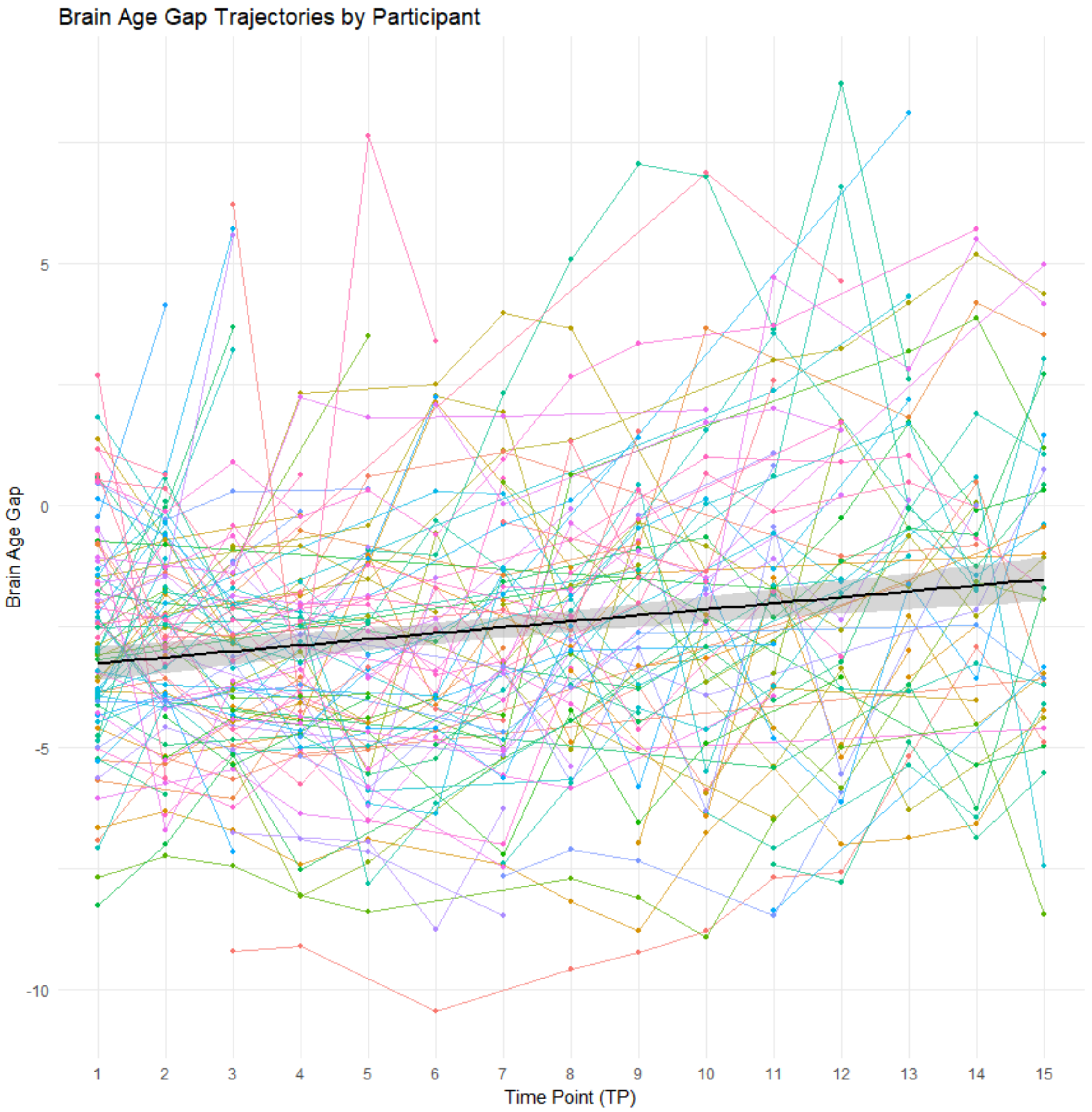


**Figure S8:** Brain-age gap x timepoint x participant trajectories across ages 12-17. Regression line of best fit with 95% confidence intervals (grey shading).

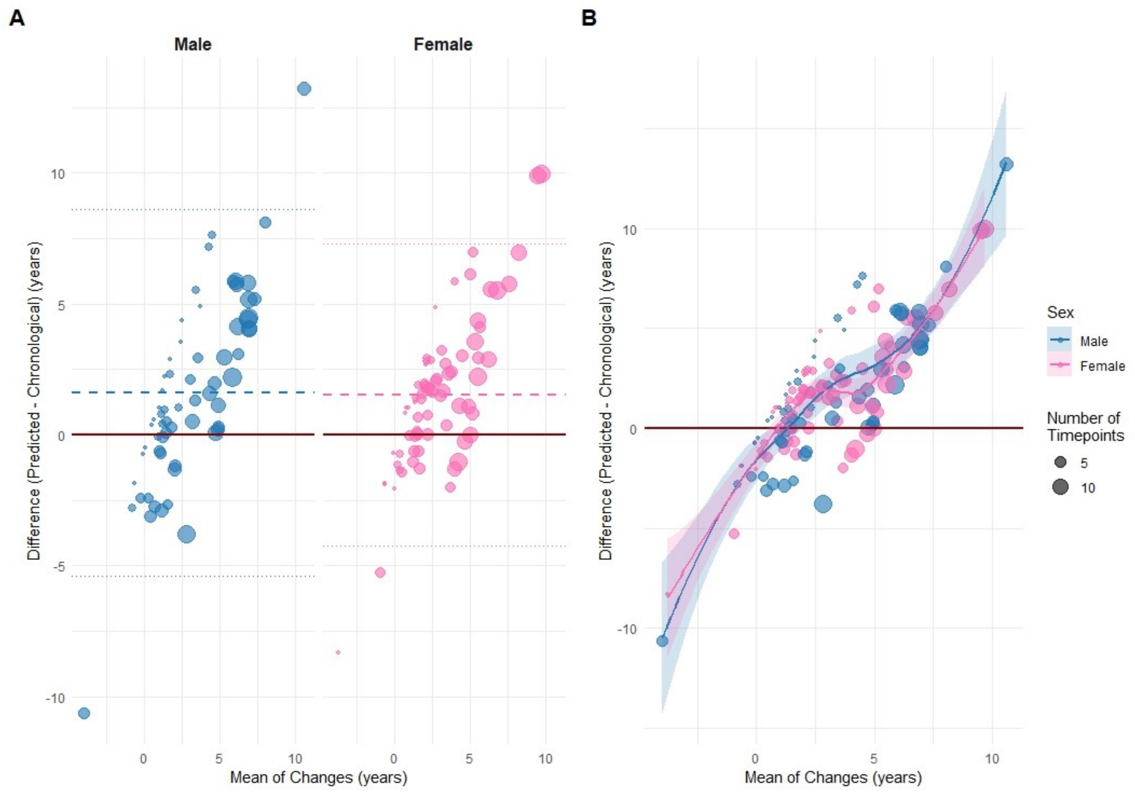


**Figure S9:** Bland-Altman Plots for change in chronological age compared with change in brain-age across ages 12-17. Size of dot reflects number of timepoints contributed by individuals to the dataset. Red reference line indicates perfect agreement. (A) Similar systematic overestimation bias was observed for males and females. Dashed line indicates actual agreement; faded lines indicate 95% of participants showing residuals between –4.78 to 7.91 years (female: -4.24 to 7.28; male: -5.39 to 8.63). (B) Dots and lines showing how the difference changes as mean increases, coloured by sex.

**Table S4: Intraclass Correlation Coefficient (ICC) estimates between all pairs of consecutive timepoints**

| Comparison | N_Paired | ICC_Pred | ICC_Gap | MAD_Unadjusted | MAD_Adjusted | LME_Slope | LME_P |
|---|---|---|---|---|---|---|---|
| TP1 vs TP2 | 73 | 0.55 | 0.576 | 1.079 | 1.079 | -3.387 | 0.422 |
| TP2 vs TP3 | 51 | 0.668 | 0.697 | 0.963 | 0.765 | -1.301 | 0.811 |
| TP3 vs TP4 | 47 | 0.669 | 0.664 | 0.76 | 0.639 | 2.942 | 0.566 |
| TP4 vs TP5 | 43 | 0.565 | 0.569 | 0.852 | 0.909 | 3.94 | 0.478 |
| TP5 vs TP6 | 23 | 0.717 | 0.725 | 1.408 | 1.667 | -14.668 | 0.057 |
| TP6 vs TP7 | 23 | 0.673 | 0.697 | 1.459 | 1.26 | -3.845 | 0.558 |
| TP7 vs TP8 | 32 | 0.653 | 0.674 | 1.759 | 1.63 | -1.078 | 0.627 |
| TP8 vs TP9 | 29 | 0.822 | 0.835 | 1.025 | 1.126 | 0.223 | 0.900 |
| TP9 vs TP10 | 21 | 0.765 | 0.769 | 1.868 | 1.688 | 2.794 | 0.704 |
| TP10 vs TP11 | 25 | 0.672 | 0.686 | 1.296 | 1.634 | -4.759 | 0.591 |
| TP11 vs TP12 | 24 | 0.748 | 0.757 | 1.251 | 0.907 | -9.703 | 0.170 |
| TP12 vs TP13 | 20 | 0.693 | 0.702 | 1.461 | 1.535 | -8.872 | 0.379 |
| TP13 vs TP14 | 24 | 0.854 | 0.854 | 1.043 | 0.893 | 8.128 | 0.082 |
| TP14 vs TP15 | 27 | 0.659 | 0.668 | 1.597 | 1.331 | 3.628 | 0.650 |

Only participants who contributed to both timepoints included in each ICC. ICC_Pred: Intraclass correlation coefficient - predicted age (the reliability/consistency of predicted BA values across two consecutive TPs), ICC_GAP: Intraclass correlation coefficient – brain age gap, MAD_Unadj: Mean Absolute Deviation – Unadjusted, MAD_Adj: Mean Absolute Deviation – Adjusted for the passage of chronological time between measurements, LME_Slope: Change (years) in BA gap between TPs, LME_P: Statistical significance.

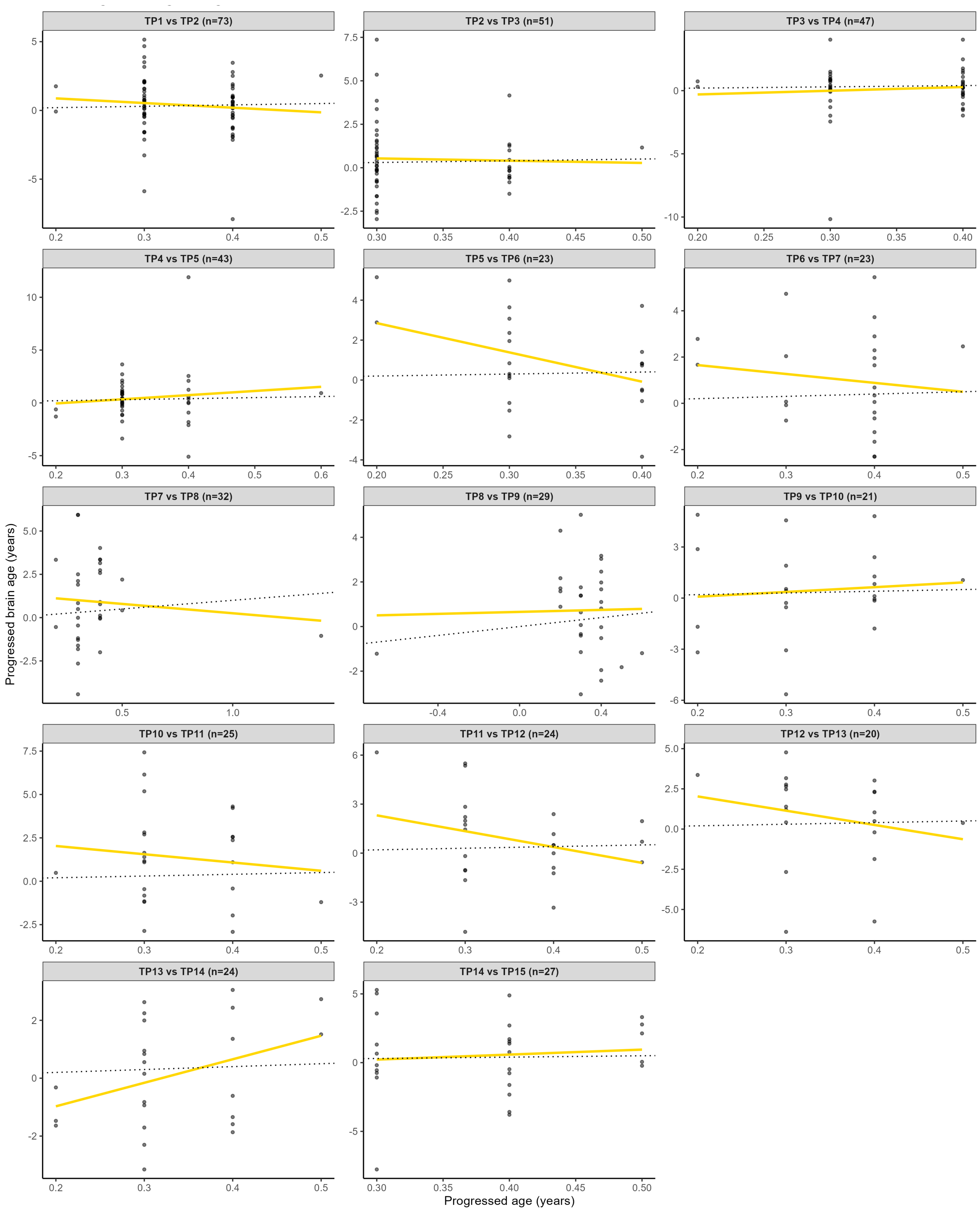


**Figure S10:** Visual representation of the Intraclass Correlation Coefficient (ICC) estimates between all pairs of consecutive timepoints

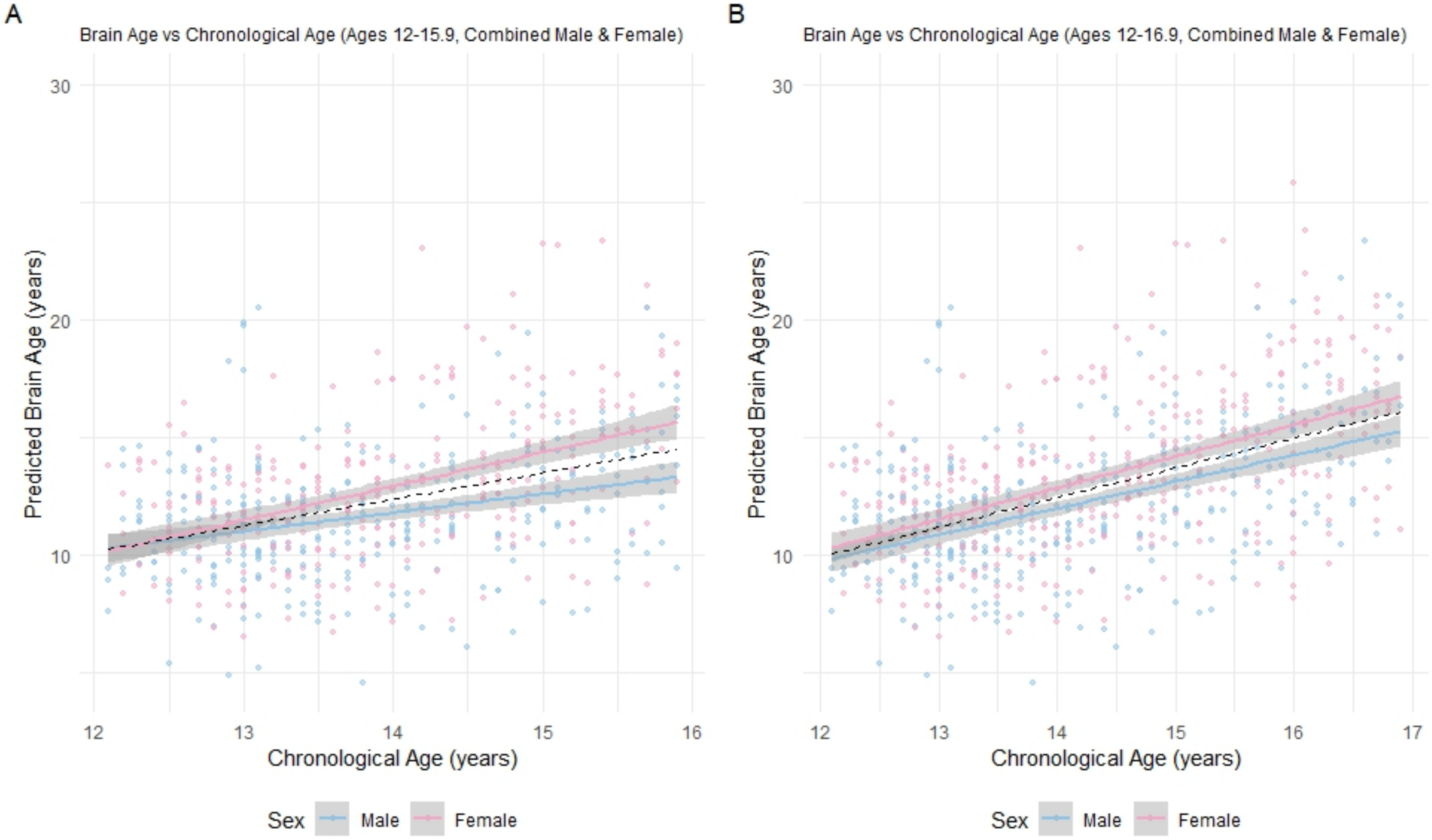


**Figure S11:** Brain-age x chronological age x sex. Regression lines of best fit indicated at the group level (red dashed line), and by sex with 95% confidence intervals (shaded). (A) Age-restricted (12-15.9 years); (B) Age-restricted (12-16.9 years); (C) Full cohort (12-17.9 years).

**Table S5: Linear mixed-effects model results across the full dataset and two restricted age ranges**

| Measure | 12-15.9 years | 12-16.9 years | 12-17.9 years |
|---|---|---|---|
| Sample size | N=126, 577 obs | N=136, 690 obs | N=138, 752 obs |
| REML criterion | 2532.25 | 3075.99 | 3397.75 |
| Age Slope (Fixed Effect) | 1.47 | 1.65 | 1.68 |
| Age Slope t-value | 16.95 | 24.79 | 28.94 |
| ID Variance (Random Effect) | 4.61 | 5.22 | 5.43 |
| ID Std. Dev. | 2.15 | 2.28 | 2.33 |
| Residual Variance | 3.04 | 3.34 | 3.64 |
| Residual Std. Dev. | 1.74 | 1.83 | 1.91 |
| Intercept | -8.02 | -10.76 | -11.13 |
